# *In vitro* monitoring of *Babesia microti* infection dynamics in whole blood microenvironments

**DOI:** 10.1101/2025.05.07.652765

**Authors:** Chao Li, Emily G. Bache, Amy L. Apgar, Danielle M. Tufts, Tagbo H.R. Niepa

## Abstract

Babesiosis – a globally emerging tick-borne infectious disease primarily caused by the intraerythrocytic piroplasm parasite, *Babesia microti* – has traditionally been studied using animal models such as mice. Compared to animal models, microfluidic-based models offer advantages, including direct analysis of human samples (such as patient blood), enhanced assay capacity (including physical/optical access, consistency, and throughput), low costs, and easy adoption. Here, we report an open microfluidic platform named “μ-Blood” for monitoring *B. microti* infection dynamics *in vitro*. Compared to other microfluidic-based models, μ-Blood allows direct examination of infected and uninfected whole blood without preprocessing steps like blood dilution or cell isolation, minimizing observer artifacts and preserving the natural whole blood microenvironment. The system enables extended (days-long) monitoring of infection dynamics, including parasite identification, parasitemia measurement, and parasite-host cell interactions, using label-free phase contrast and fluorescence confocal microscopy. With its open microfluidic configuration, μ-Blood provides an *in vitro* model for studying blood-borne infections while maintaining integrity of the whole blood microenvironment.

## INTRODUCTION

*Babesia microti* – an intraerythrocytic protozoan parasite from the Apicomplexa phylum – infects and damages red blood cells (RBCs) in animal and human hosts, posing a significant health burden, particularly in immunocompromised individuals (*1, 2*). Despite certain advances in knowledge, cases of babesiosis continue to rise annually (*3*) due to the lack of full understanding of the unconventional host cycles of the parasite, pathogenesis, parasite resistance and relapse against treatment, and emergence of new mutant species (*4–6*). For example, *B. microti* can be transmitted horizontally from tick to host, vertically from mother to offspring, or via blood transfusion or organ donation (*7–9*). Distinct from other erythrocytic parasites, *B. microti* exploits host cycles of different lengths to gain survival advantage in hostile environments (*4*). However, the mechanisms employed by the parasite during transmission and infection are poorly understood.

The development of new strategies for infection intervention, including prevention and treatment, requires a deeper understanding of infection dynamics and the biophysical and chemical mechanisms of parasites within host environments (*4*). Animal models, such as mice, have been the gold standard for studying the mechanisms of parasite infection (*10, 11*). Specifically, in human-infecting *Babesia* spp studies, severe combined immunodeficiency (SCID) mice (*12*) are often used to maintain an infected colony, as these mice lack a functional immune system. However, in animal models, real-time parasite-host cell interactions, direct physical and optical (i.e., imaging) access of the parasites are inherently limited. To monitor and study infection dynamics, without sacrificing the host, small blood volumes are collected from mice (10% mouse total blood volume every two weeks) and then analyzed *in vitro* using blood smears (*13*), serology (*14*), and/or polymerase chain reaction (PCR) analyses (*15*). These currently adopted *in vitro* analysis methods provide only a “snapshot in time” of infection dynamics at cellular or molecular levels (e.g., antibody, DNA) at a given time point or through several time points discretely. In addition, the use of mouse models is constrained by the need for specialized facilities (e.g., mouse laboratories), extensive personnel training, and ongoing maintenance fees (*10, 11*).

The limitations discussed above are rooted in poor cultivability of the parasite outside the host system (*16, 17*). Once leaving the host, the parasite exhibits low levels of proliferation and propagation due to the altered microenvironment from *in vivo* to *in vitro*. To mitigate the use of lab animals and enable continuous monitoring of infection dynamics *in vitro*, blood cell functional assays, i.e., interrogation of the parasite-host cell interactions over a period (e.g., hours, days, or longer), are needed. Ideally, the blood cell functional assay platform should allow i) continuous (days-long) cultivation of the parasite in an *in-vivo*-like microenvironment, ii) spatiotemporal sample manipulation, and iii) continuous monitoring of the infection dynamics *in situ* with real-time readouts.

Recently, we developed an open microfluidic whole blood cell functional assay platform known as μ-Blood (Fig. 1, Fig. S1) (*18*). μ-Blood is established upon an Exclusive Liquid Repellency (ELR, see Fig. S2) (*19, 20*)-Under-oil Open Microfluidic System (UOMS) (*21*). Compared to other microfluidic blood assay platforms, μ-Blood features: i) direct use of unprocessed whole blood for both input and through an assay (days long) preserving the original host-specific whole blood microenvironment (*18*), ii) streamlined operation steps (i.e., removal of blood dilution or cell isolation) and reduced observer artifacts (i.e., blood sample contamination, loss of the inherent signaling molecules in blood) (*22*), iii) free physical access to samples on the device with minimal system disturbance (e.g., particle/bubble clogging commonly seen in closed-system microfluidics, irreversible device damage from physical access) (*23*), iv) autonomously regulated oxygen microenvironment (AROM) (*24*), v) various/combined optical access (i.e., phase contrast (*23*), epifluorescence/confocal (*25*), multiphoton (*26*), Raman (*27*), and infrared (IR) (*28*) microscopy), vi) alignment with open-system standard and lab automation (*29*), and vii) low adoption/implementation barriers.

**Fig. 1.**
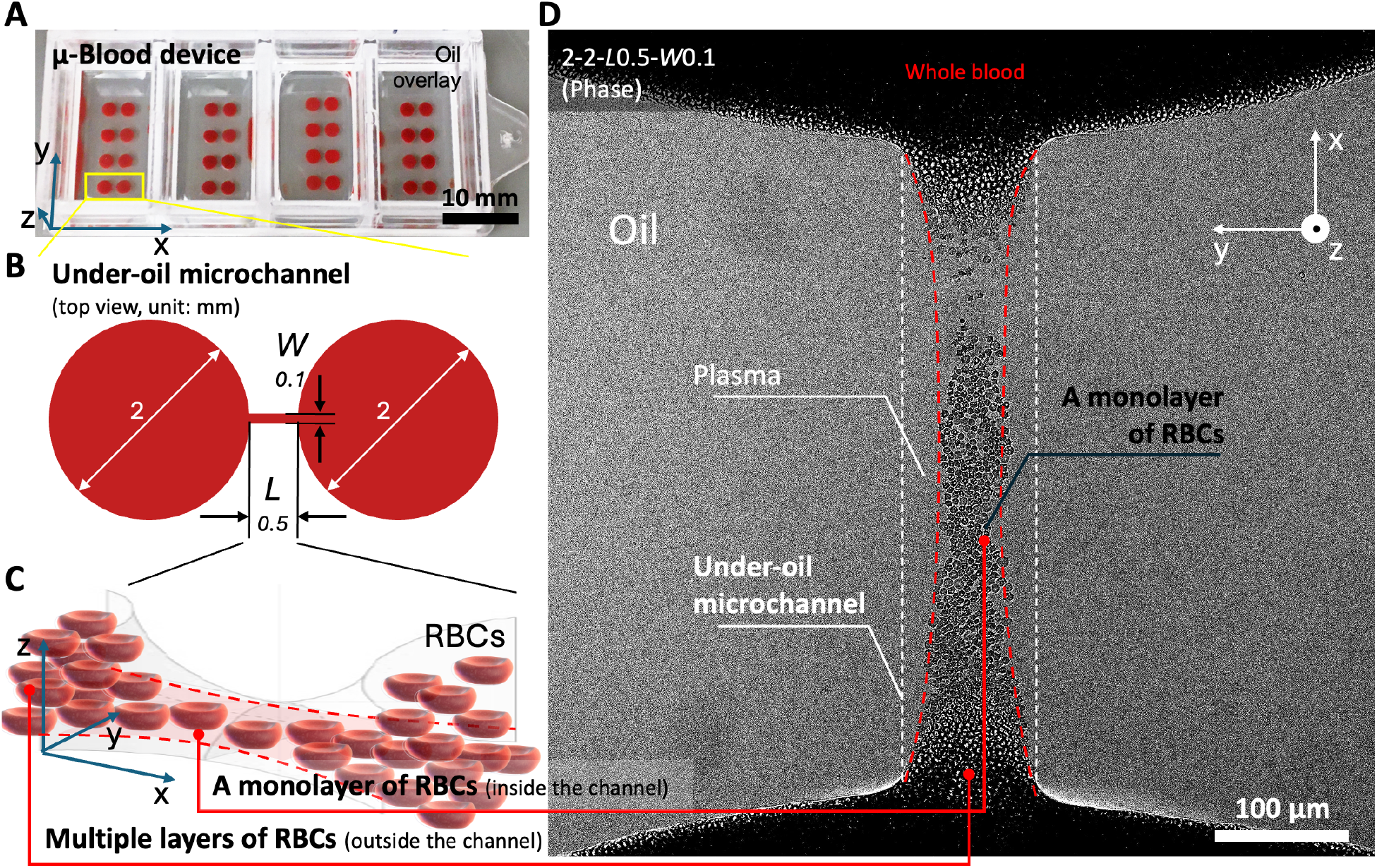
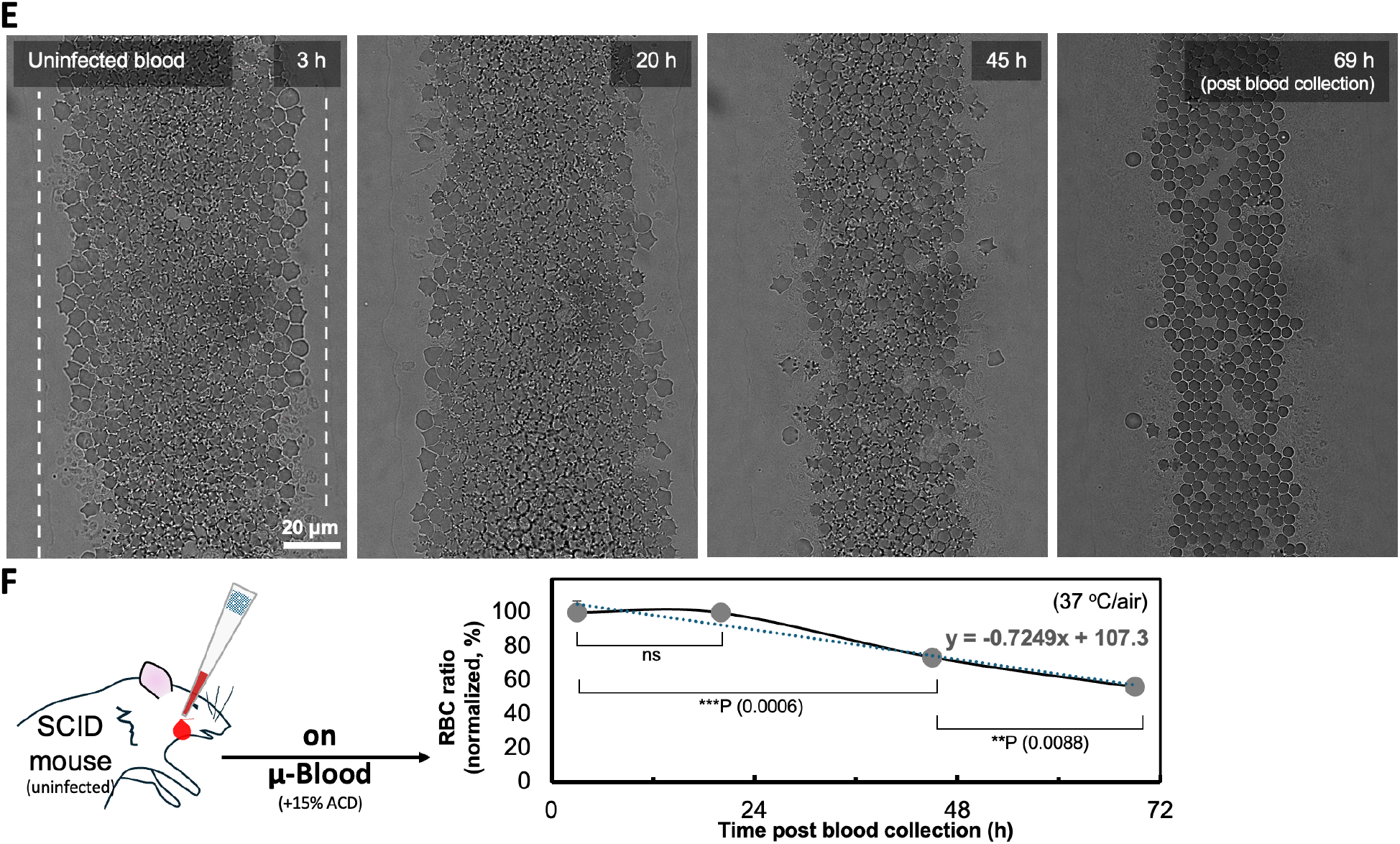
μ-Blood device settings and the under-oil whole blood assay microenvironment. (**A**) Image of the μ-Blood device fabricated with Lab Tek II 4-well chambered coverglass (6 cm × 2.4 cm, the coverglass bottom – 0.15 mm in thickness) (Fig. S1C). The device holds a 4 × 4 array of under-oil microchannels (highlighted by the yellow box). The oil overlay is Fluorinert FC-40 (500 μL per well). The xyz coordinates indicate the orientation of microchannels on the μ-Blood device. (**B**) Two-dimensional (2D) schematic (top view) shows the dimensions of the under-oil microchannels, which are 0.5 mm in length (*L*), 0.1 mm in width (*W*), connecting two circular spots 2 mm in diameter, written as 2-2-*L*0.5-*W*0.1. (**C**) Three-dimensional (3D) schematic shows the hyperbolic paraboloid shape of the under-oil microchannels. Particles (e.g., RBCs – about 6-8 μm in diameter, 2 μm in thickness) are excluded from the areas if the media layer thickness is less than the minimal size of the particles (e.g., the thickness of RBC). In the under-oil microchannels, RBCs spontaneously form a monolayer denoted by the red dashed lines. (**D**) Microscopic image [confocal – phase, objective – 25× (water immersion), top view] of an under-oil microchannel (white dashed lines) loaded with whole blood (from an SCID mouse). The multiple layers of RBCs outside the channel block optical illumination through blood. By contrast, the monolayer of RBCs (red dashed lines) inside the channel allows uncompromised optical access to the cells in preserved whole blood microenvironment. The image orientation on the microscope computer screen is rotated 90° counterclockwise compared to the orientation of the μ-Blood device shown in (**A**). (**E**) Images (phase) and (**F**) plot show mouse RBC (uninfected with 15% ACD) stability on μ-Blood. The dotted line shows the linear fitting (R^2^ = 0.9436). Error bars, mean ± s.d. **P ≤ 0.01, ***P ≤ 0.001, and ns – not significant.

In this study, we utilized μ-Blood to interrogate whole blood collected from *B. microti*-infected SCID mice. We validated the experimental protocols and quantitative monitoring of the infection dynamics over 4 days *in vitro*. Specifically, μ-Blood supports i) label-free (phase contrast-based) identification of *B. microti* in whole blood, ii) parasitemia measurement with improved accuracy, and iii) days-long monitoring of the dynamics of *B. microti*-infected RBCs. These results demonstrate the ability of μ-Blood for *in vitro* studying blood-borne infections with donor-specific, preserved whole blood microenvironments.

## RESULTS

### Whole blood assay microenvironment on μ-Blood

This study is a new implementation of μ-Blood (Fig. 1) to investigate an intracellular parasitic pathogen and infection dynamics (i.e., progression of parasite behavior over time). μ-Blood can be fabricated on a broad range of surfaces including silicon, glass, and plastics (e.g., polystyrene, polycarbonate) (Fig. S1C-I). Here we chose Lab Tek II chambered coverglass as the device carrier to leverage its standardization and compatibility with the microscopy systems (Fig. 1A, Fig. S1C-III). Specifically, we adopted a 4-well chambered coverglass to hold a 4 × 4 array of under-oil microchannels. Each well houses four microchannels with wells physically separated from each other to minimize cross-sample contamination (Fig. 1A). The under-oil microchannels were designed to be 0.5 mm in length (*L*) and 0.1 mm in width (*W*) connecting the two circular spots 2 mm in diameter (written as 2-2-*L*0.5-*W*0.1) (Fig. 1B).

Defined by the physics of oil-media interface, the under-oil microchannels take a hyperbolic paraboloid shape (Fig. 1C) with a liquid-liquid interface ceiling. This microchannel geometry is inherently distinct from the engineered microfluidic channels that commonly take a constant ceiling height. Known from previous studies, a 0.1 mm channel width leads to an averaged media layer thickness of <3 μm in the under-oil microchannel (*18*). Whole blood collected from a *B. microti-*infected SCID mouse (Fig. S1A) was loaded into the microchannels via the “under-oil sweep distribution” technique (Fig. S1C-III) (*20, 21*). Briefly, a hanging drop of whole blood (10 to 20 μL) was dragged across the patterned surface under oil using a large orifice pipette tip (2 mm in diameter at the end opening). Directed by the double-ELR [i.e., under-oil water ELR (the unpatterned areas) + under-water oil ELR (the patterned areas)] physics (*20*), whole blood was spontaneously and exclusively distributed to the patterned areas, leaving the unpatterned areas free from blood dispensing and biofouling (Fig. 1 A and D, Fig. S2). Defined by the hyperbolic paraboloid geometry, a monolayer of RBCs spontaneously formed in a microchannel, excluding the areas along the edges in the microchannel (Fig. 1D, the areas occluded between the white and adjacent red dashed lines).

RBCs are dense in whole blood (4-7 million per μL of blood). In addition, the hemoglobin (Hb) in RBCs strongly absorbs visible light. Together, these factors lead to poor optical access (especially with phase contrast) through multiple layers of RBCs (Fig. 1D, the blood cells outside the channel causing the area to appear dark due to the high cell density and limited light transmission). By contrast, the monolayer of RBCs in the microchannel allowed for uncompromised optical access to the cells (Fig. 1D, the blood cells inside the channel between the red dashed lines) without the need for blood preprocessing (e.g., blood dilution or cell isolation) or fluorescence tagging. The monolayer of RBCs enables label-free monitoring of living cells directly in unprocessed whole blood with minimized observer artifacts. On μ-Blood, we monitored the change of uninfected mouse RBCs in 3 days as the assay baseline (Fig. 1E). The results showed that mouse RBCs without infection (supplemented with 15% ACD for anticoagulation) were relatively stable in 24 h and then progressively degraded *in vitro* (37 °C/air) at a rate about 22% per day (Fig. 1F).

### Improved identification accuracy of *B. microti* in whole blood

We first attempted to identify *B. microti* in whole blood by testing the capability of label-free imaging, specifically phase contrast-based parasite identification on μ-Blood. MitoTracker Green (an irreversible mitochondrial dye that stains both live and dead mitochondria) was used as a reference for *B. microti* identification (Fig. 2A). Given the fact that RBCs lack mitochondria, MitoTracker Green selectively stains *B. microti* (with one mitochondrion per *B. microti* cell or two mitochondria co-existing in a dividing *B. microti* cell) in a whole blood sample.

**Fig. 2.**
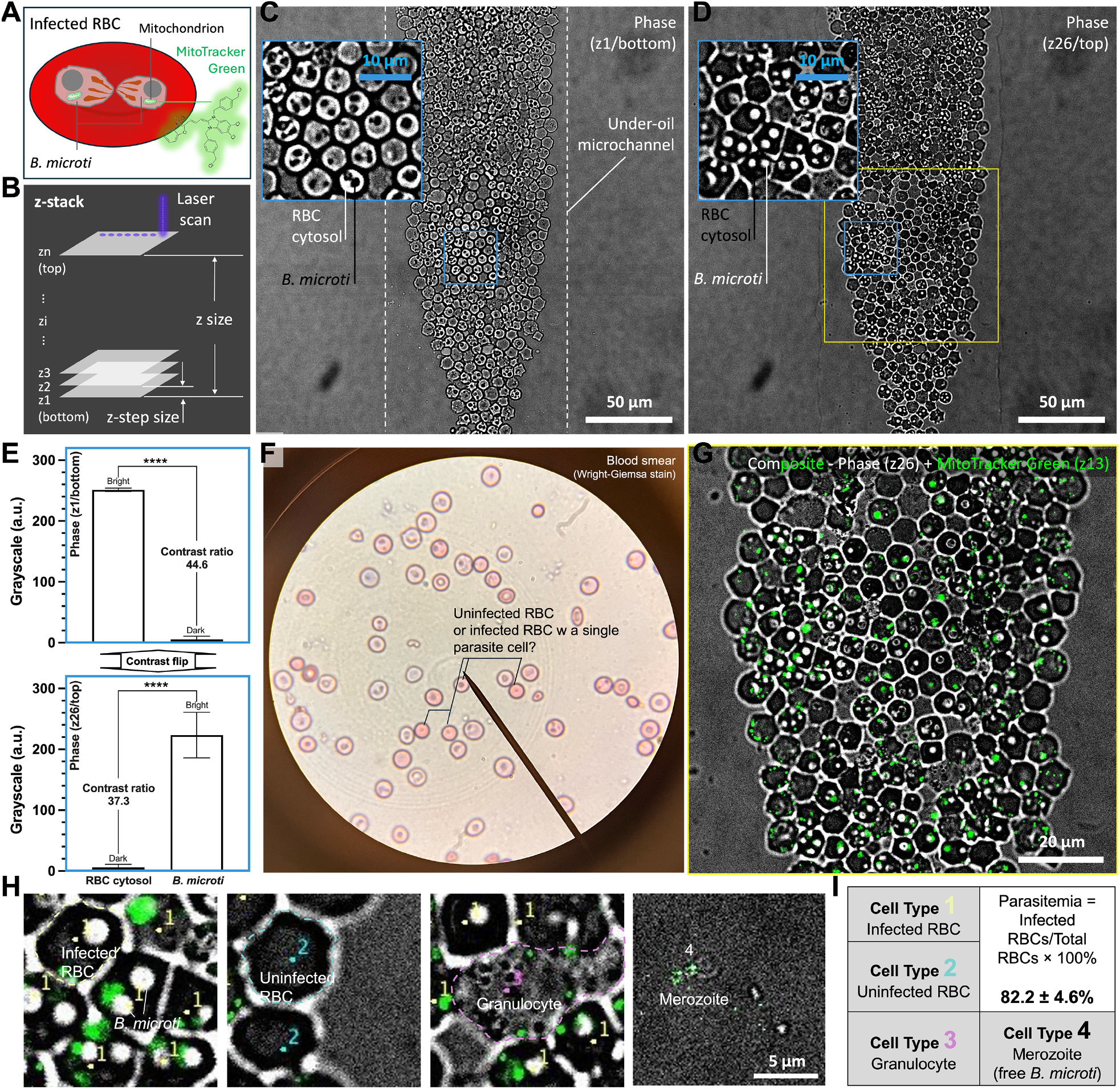
Identification of *B. microti* in whole blood and parasitemia measurement with confocal imaging. (**A**) Schematic shows the intraerythrocytic parasites and staining strategy. MitoTracker Green [Excitation/Emission (Ex/Em) 490 nm/516 nm] was used as the reference to distinguish *B. microti* (with mitochondrion) from RBC (without mitochondrion). (**B**) z-stack setting of the confocal imaging. The scan was set from bottom (z1) to top (zn) of the monolayer of RBCs in the under-oil microchannel, with z-step size set to 0.2 μm through 26 slices with a frame rate of 0.024/s (i.e., 41.7 s per frame, the total time of scan – 18 min). (**C**) Phase image [8-bit, objective – 63× (oil immersion)] of the bottom slice (z1) shows RBC cytosol in bright pixels and *B. microti* cells in dark pixels. The white dashed lines denote the boundary of the under-oil microchannel. (**D**) Phase image [8-bit, objective – 63× (oil immersion)] of the top slice (z26) shows RBC cytosol in dark pixels and *B. microti* cells in bright pixels. Insets in (**C**) and (**D**) show the zoomed-in images with a region of interest (ROI) of 30 μm × 30 μm (the blue solid-line box). (**E**) Phase contrast flipped between the bottom slice and the top slice (Movie S1). RBC cytosol (n = 14); *B. microti* (n = 18). Error bars, mean ± s.d. ****P ≤ 0.0001. (**F**) Blood smear image shows the RBCs with Wright-Giemsa stain (blue-ish color). (**G**) Composite image (phase-z26 + MitoTracker Green-z13) of the ROI of 100 μm × 100 μm (the yellow solid-line box) in (**D**) shows stained mitochondria in *B. microti-*infected RBCs. (**H**) Cell types identified from cell count in (**G**) including 1. Infected RBC, 2. Uninfected RBC, 3. Granulocyte, and 4. merozoite (i.e., free *B. microti*). (**I**) Cell counts (replicates ×3) of each cell type (Fig. S3C) and parasitemia calculation (Table S1). The sample in this experiment was whole blood from a *B. microti*-infected SCID mouse. The images were recorded on μ-Blood at 4 h post blood collection with the device incubated at 37 °C in ambient air.

Targeting the monolayer of RBCs in the microchannel (Fig. 1D), we created a three-dimensional z-stack from bottom to top of the monolayer with a 63× oil immersion objective and two channels including phase contrast and MitoTracker Green (Fig. 2A). The z-stack was set based on the MitoTracker Green channel, scanning 5.2 μm z size through the monolayer of RBCs including 26 steps (or slices) from z1 to z26 with 0.2 μm z-step size (Fig. 2B, Movie S1). The fluorescence intensity of MitoTracker Green increased from zero at the bottom slice (z1), maximized at the middle slice (z13), and then dropped back to zero at the top slice (z26) (Fig. S3A). In comparison, phase contrast flipped between the bottom slice and the top slice (Movie S1). The phase contrast channel showed maximum grayscale (8-bit, 0-255) contrast ratio between RBC cytosol and *B. microti* cells at the bottom slice (contrast ratio – 44.6, RBC cytosol – bright, *B. microti* cell – dark) and the top slice (contrast ratio – 37.3, RBC cytosol – dark, *B. microti* cell – bright) [RBC cytosol (sample size n = 14), *B. microti* (n = 18), P ≤ 0.0001, t-test] (Fig. 2 C to E). At the middle slice (z13) where the fluorescence intensity of MitoTracker Green was maximized, the grayscale contrast ratio dropped to ∼1.4 (Fig. S3B), only ∼4% of the contrast ratio compared to the top or bottom slice (Fig. 2E).

To identify *B. microti*-infected RBCs (Fig. 2G), we combined the middle slice (z13) of MitoTracker Green (Fig. S3A) and the top slice (z26) of phase contrast (Fig. 2D). Because of the high contrast in grayscale between the bottom and top slices (Fig. 2 C to E), we chose to use the top slice of phase contrast to also highlight the RBC cytosol (darker due to Hb) and parasites [brighter (Fig. 2D, inset) + maximum fluorescence intensity (Fig. S3A)]. Using the combined images on μ-Blood (Fig. 2G), we were able to identify four cell types: 1. Infected RBC, 2. Uninfected RBC, 3. Granulocyte, and 4. Merozoite (i.e., free *B. microti*) (Fig. 2H). Cell Type 1 showed an RBC diameter of 6-8 μm with a dark cytosol, intracellular *B. microti* cells visualized by bright dots or rings with a typical diameter of 1-2 μm, and green fluorescence of mitochondria. Cell Type 2 showed similar RBC dimensions to Cell Type 1, but with no intracellular *B. microti* cells or green fluorescence of mitochondria. Cell Type 3 showed larger cell size >10 μm and lighter cytosol compared to RBCs. Also, granules were observed in most of the Cell Type 3, suggesting non-RBC cells such as granulocytes (e.g., neutrophils). SCID mice are genetically modified to lack lymphocytes (B cells, T cells, and NK cells) but still produce other white blood cells such as granulocytes (e.g., neutrophils) (*30*). Cell Type 4 showed an averaged cell size around 2-3 μm identified as free parasite cells (i.e., merozoites) released into plasma. Further characterization and analysis of the cellular objects identified in *B. microti* infection on μ-Blood are discussed in the following section.

Blood smear with Wright-Giemsa stain was performed as the current standard for parasitemia measurement (Fig. 2F). However, the contrast between the uninfected and infected RBCs with Wright-Giemsa stain appeared low, limiting the measurement accuracy and consistency (Table S1). In comparison, confocal imaging on μ-Blood clearly established a parasitemia level of 82.2 ± 4.6 in the blood samples (Fig. 2 H and I), highlighting an improved accuracy and consistency in parasitemia measurement using the platform. While MitoTracker Green was primarily used in μ-Blood to confirm and identify *B. microti*, parasitemia measurement on μ-Blood can be completed label free (using phase contrast), without any fluorescence tagging. Label-free parasitemia measurement reduces operation steps, time, costs, and observer artifacts.

### Monitoring of parasite viability using MitoTracker

We further demonstrated the capability of monitoring and evaluating parasite viability in whole blood on μ-Blood. Cell viability can be evaluated using different types of fluorescence dye, typically by visualizing the integrity of the cytoplasm/nuclear membrane [i.e., various membrane-impermeable DNA stains such as propidium iodide (PI)] (*31*), enzyme level in cytosol [i.e., esterase level indicated by conversion of Calcein AM (nonfluorescent) to Calcein (fluorescent)] (*32*), or mitochondrial activity (*33*). DNA stains are known as cytotoxic and typically used for end-point viability assay only. Calcein AM gets hydrolyzed in plasma due to the esterases (e.g., albumin esterase, butyrylcholinesterase (BChE), and paraoxonase (PON1)] in plasma. Hydrolysis of Calcein AM in plasma contributes to a high fluorescence background, interfering with the signals from cells (Fig. S4 A and B). While the fluorescence background can be lowered by diluting/washing the blood cells in phosphate-buffered saline (PBS) (Fig. S4C), the dilution/wash steps alter the original whole blood microenvironment due to the dilution or removal of signaling factors in plasma.

To monitor parasite viability, we used live mitochondrial dye (MitoTracker Orange) as a probe quantifying the mitochondrial activity (Fig. 3 A and B). Like MitoTracker Green, MitoTracker Orange can be directly used in whole blood with a low working concentration of 0.1 μM, avoiding blood dilution or wash steps. We found that i) MitoTracker Orange stains live mitochondria directly in whole blood without performing any wash steps (Fig. 3A), ii) the dye is stable against high-frequency laser scan, showing no noticeable photobleaching against 2% max. laser power (Leica, Stellaris 5, WLL) and 360 scans through 6 h (Fig. 3B), iii) the cytotoxicity of the dye on mitochondria is minimal, and iv) the fluorescence intensity of dead/dying mitochondria drops significantly compared to live mitochondria (Fig. 5 J and K).

**Fig. 3.**
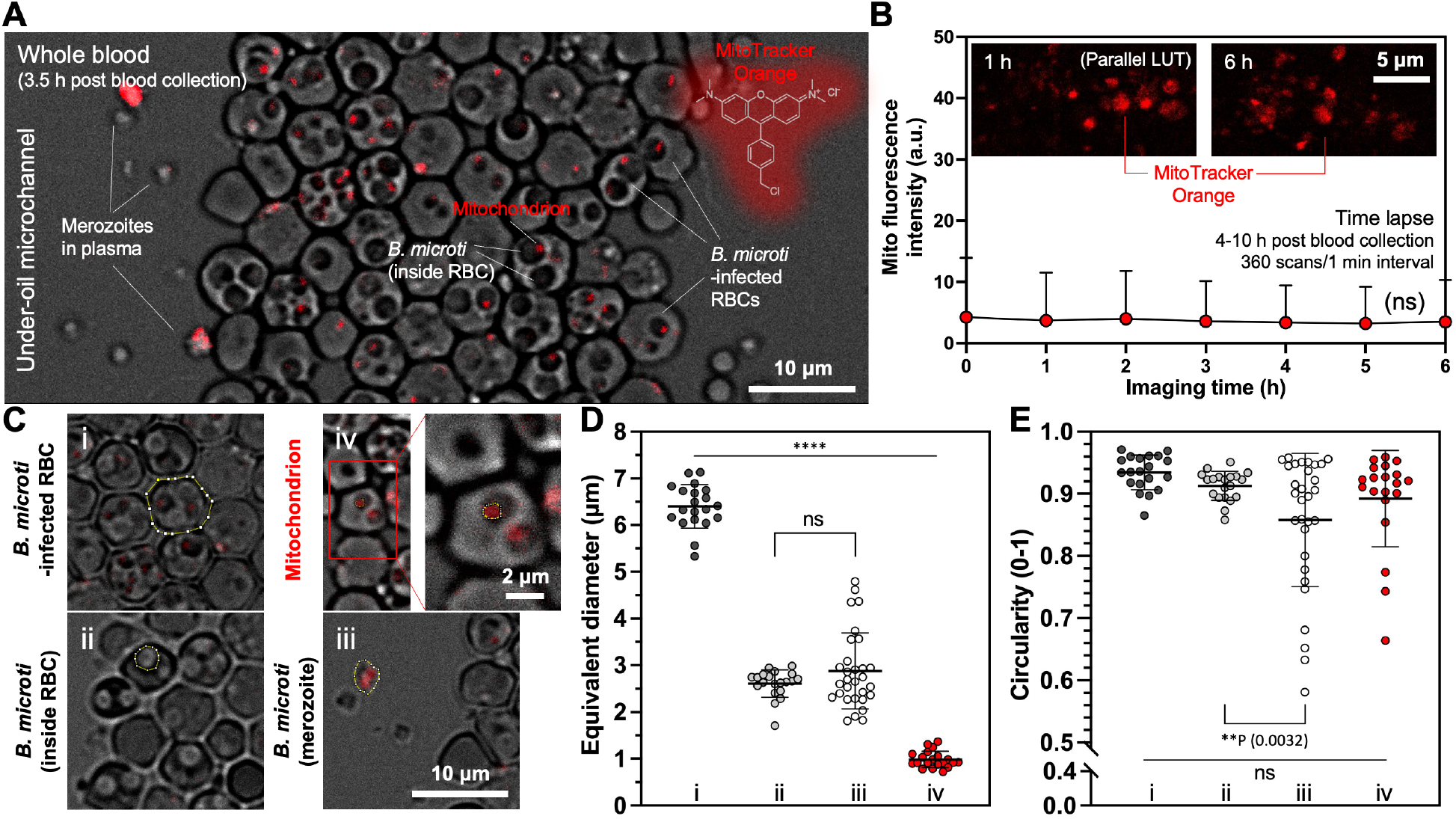
Parasite viability probed with mitochondrial activity. (**A**) Overview image [composite – phase + MitoTracker Orange (Ex/Em 554 nm/576 nm), 8-bit, objective -20× (dry)] shows an under-oil microchannel filled with whole blood from a *B. microti* infected SCID mouse. The cellular objects identified (from large to small) include *B. microti*-infected RBC, *B. microti* (inside RBC), *B. microti* (outside RBC, i.e., merozoite), and *B. microti* mitochondrion (red fluorescence). **(B)** Photostability of MitoTracker Orange. The dye was scanned 360 times through 6 h, showing no noticeable decay in fluorescence intensity. (**C**) The representative image of (i) *B. microti*-infected RBC, (ii) *B. microti* (inside RBC), (iii) *B. microti* (merozoite), and (iv) mitochondrion. The yellow-line polygon boxes denote the ROI of each type of cell in shape analysis. (**D**) Equivalent diameter [SQRT(4×surface area of ROI/π)] and (**E**) circularity (0 – least circular; 1 – perfectly circular) of the cellular objects shown in (**C**). (i) *B. microti*-infected RBC (n = 20); (ii) *B. microti* (inside RBC) (n = 20); (iii) *B. microti* (merozoite) (n = 30); (iv) mitochondrion (n = 20). Error bars, mean ± s.d. **P ≤ 0.01, ****P ≤ 0.0001, and ns – not significant.

Using phase contrast and MitoTracker Orange channels combined, we were able to distinguish and identify *B. microti*-infected RBCs (against uninfected RBCs), *B. microti* inside an RBC, free *B. microti* (i.e., merozoites) in plasma, and mitochondria in the parasite cells (Fig. 3C). *Babesia microti*-infected RBCs 3-5 h post blood collection exhibited a diameter of 6-8 μm with a circular shape (average circularity, 0.93) (Fig. 3 D and E). The parasite cells were significantly smaller (average 2.5 μm in diameter) compared to the infected RBCs [infected RBCs (n = 20), *B. microti* (inside RBC) (n = 20), *B. microti* (merozoite) (n = 30), P ≤ 0.0001, One-way ANOVA]. When inside an RBC, the parasite cells had a largely circular shape (average circularity, 0.91). By contrast, the free parasite cells (i.e., merozoites) in plasma showed significantly larger deformability (average circularity, 0.86) with the circularity as low as 0.58 [*B. microti* (inside RBC) (n = 20), *B. microti* (merozoite) (n = 30), P = 0.0032, One-way ANOVA], as is expected from the teardrop shape of *B. microti* merozoites (*5*). Mitochondria inside the parasite cells (n = 30) were ∼1 μm in diameter with circularity as low as 0.66. These significant differences among the host cell, the parasite cell, and mitochondria identified on μ-Blood lay the foundation for robust and accurate information extraction through the tracking of infection dynamics.

### Quantification of Hb level with RBC cytosol grayscale

*Babesia microti* burdens the host system by infecting and damaging RBCs (*4*). The capability of a blood assay platform to monitor *B. microti*-RBC interactions in whole blood microenvironment is essential to understand the infection dynamics. Here, we quantified the change of Hb level of RBC cytosol before and after RBC cytoplasmic membrane damage (Fig. 4).

**Fig. 4.**
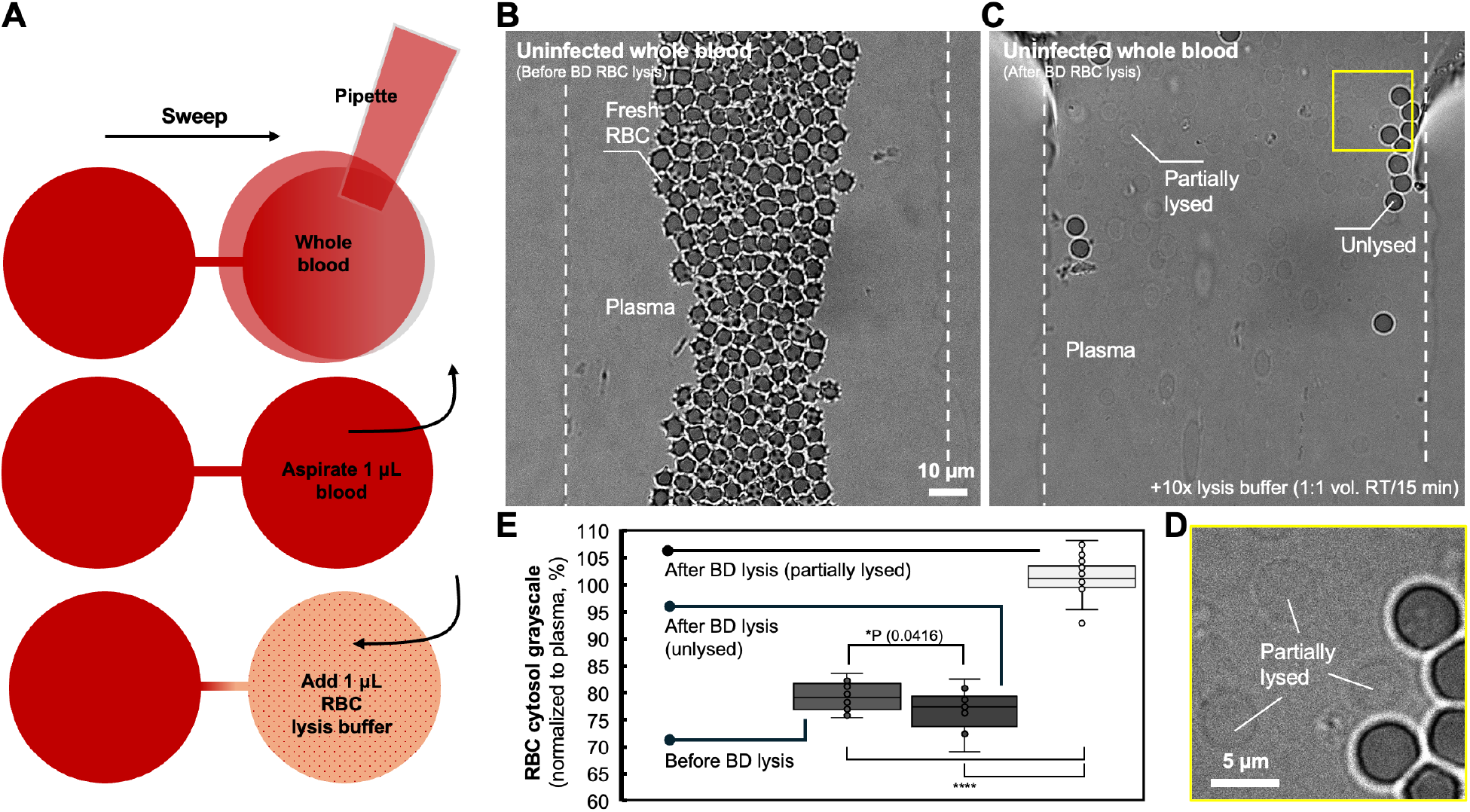
Comparison of RBC cytosol grayscale in RBC lysis. (**A**) Schematic showing the workflow of blood loading and lysis on μ-Blood. (**B**) Phase image of fresh uninfected RBCs in the under-oil microchannel (white dashed lines). **(C)** RBCs after partial lysis [BD, 10× lysis buffer, incubated at room temperature (RT) for 15 min]. The yellow box highlights the zoomed-in image (**D**) showing the cytoplasmic membrane of unlysed and partially lysed RBCs. (**E**) RBC cytosol grayscale (8-bit) normalized to plasma (n = 15) before (n = 15) and after lysis including unlysed (n = 12) and partially lysed (with lighter cytosol and visible cytoplasmic membrane, n = 20). Error bars, mean ± s.d. *P ≤ 0.05 and ****P ≤ 0.0001.

RBCs can be fully or partially lysed using RBC lysis buffers. On μ-Blood, we loaded uninfected blood from an SCID mouse into under-oil microchannels with sweep distribution (Fig. 4A). The RBC lysis buffer was added to one end of the microchannel, triggering partial RBC lysis (Fig. 4 B to D). RBC cytosol grayscale was analyzed and compared before and after lysis (Fig. 4E). Due to Hb absorption of visible light, the cytosol of fresh uninfected RBCS (n = 15) is significantly darker than plasma (n = 15, P ≤ 0.0001, t-test) on phase contrast. After lysis buffer treatment, fully lysed RBC cells were completely gone, merging with plasma (Fig. 4C). Partially lysed RBC cells (n = 20) showed significantly lighter cytosol compared to fresh uninfected RBCs (n = 15, P ≤ 0.0001, t-test), with still visible cytoplasmic membrane (Fig. 4D). A small group of RBCs were left unlysed (n = 12) from the lysis buffer treatment, with a slightly darker cytosol compared to the RBCs before lysis (n = 15, P = 0.0416, t-test) (Fig. 4D). These results provide the basis for monitoring the Hb level change of *B. microti*-infected RBCs on μ-Blood through an infection process.

### Dynamics of *B. microti*-infected RBCs monitored in real time on μ-Blood

Last, we monitored the dynamics of *B. microti*-infected RBCs through a 4-day time course on μ-Blood with the label-free RBC cytosol grayscale protocol and the MitoTracker Orange-based parasite viability probe established above. In this study, the earliest imaging was done 3-4 h post blood collection, determined by logistics including blood sample collection, transport, and device preparation (Fig. S1A). On μ-Blood, two types of *B. microti*-infected RBCs were identified, including RBCs with initial Hb and RBCs with low Hb (Fig. 5 A and B), determined using phase contrast microscopy.

**Fig. 5.**
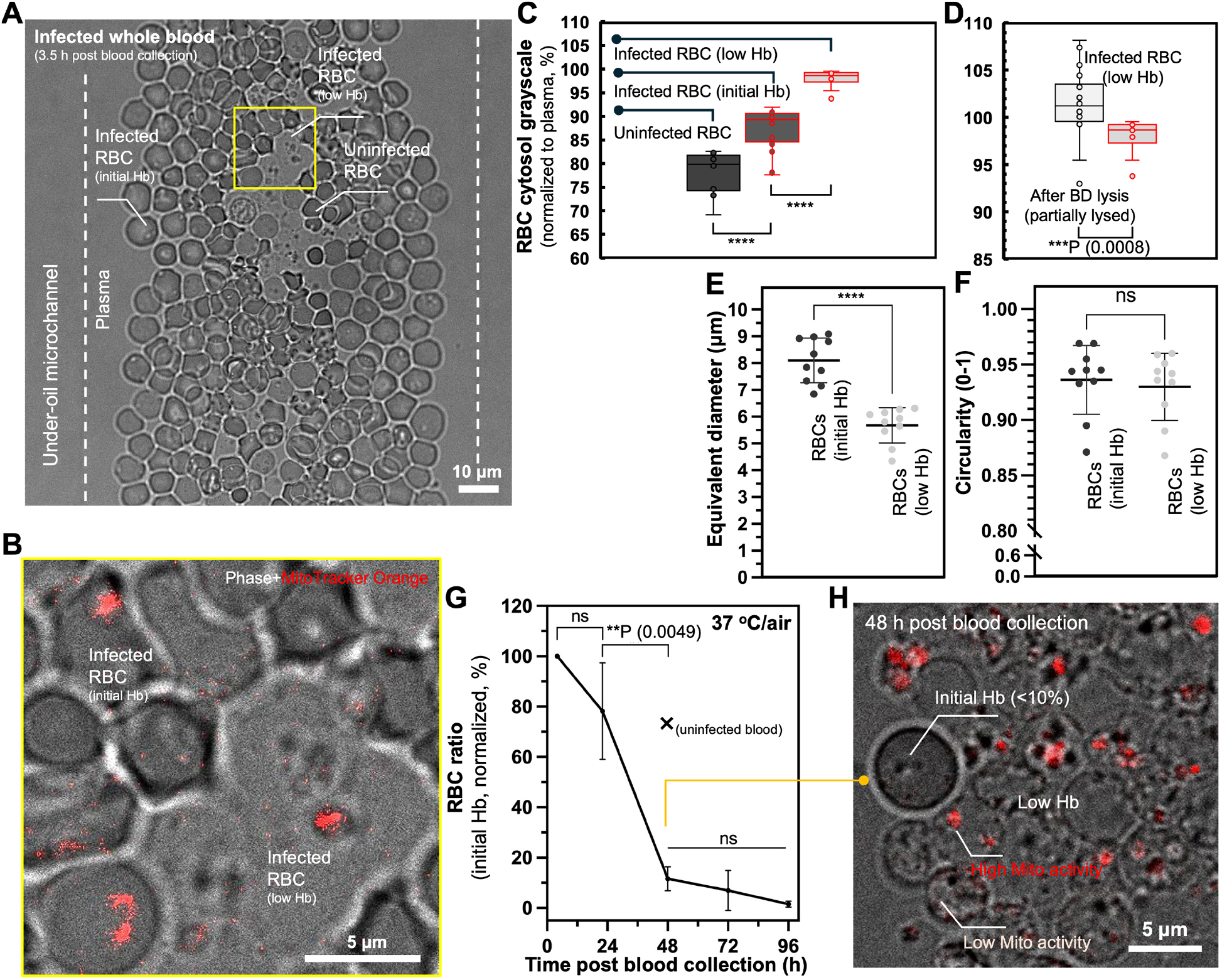
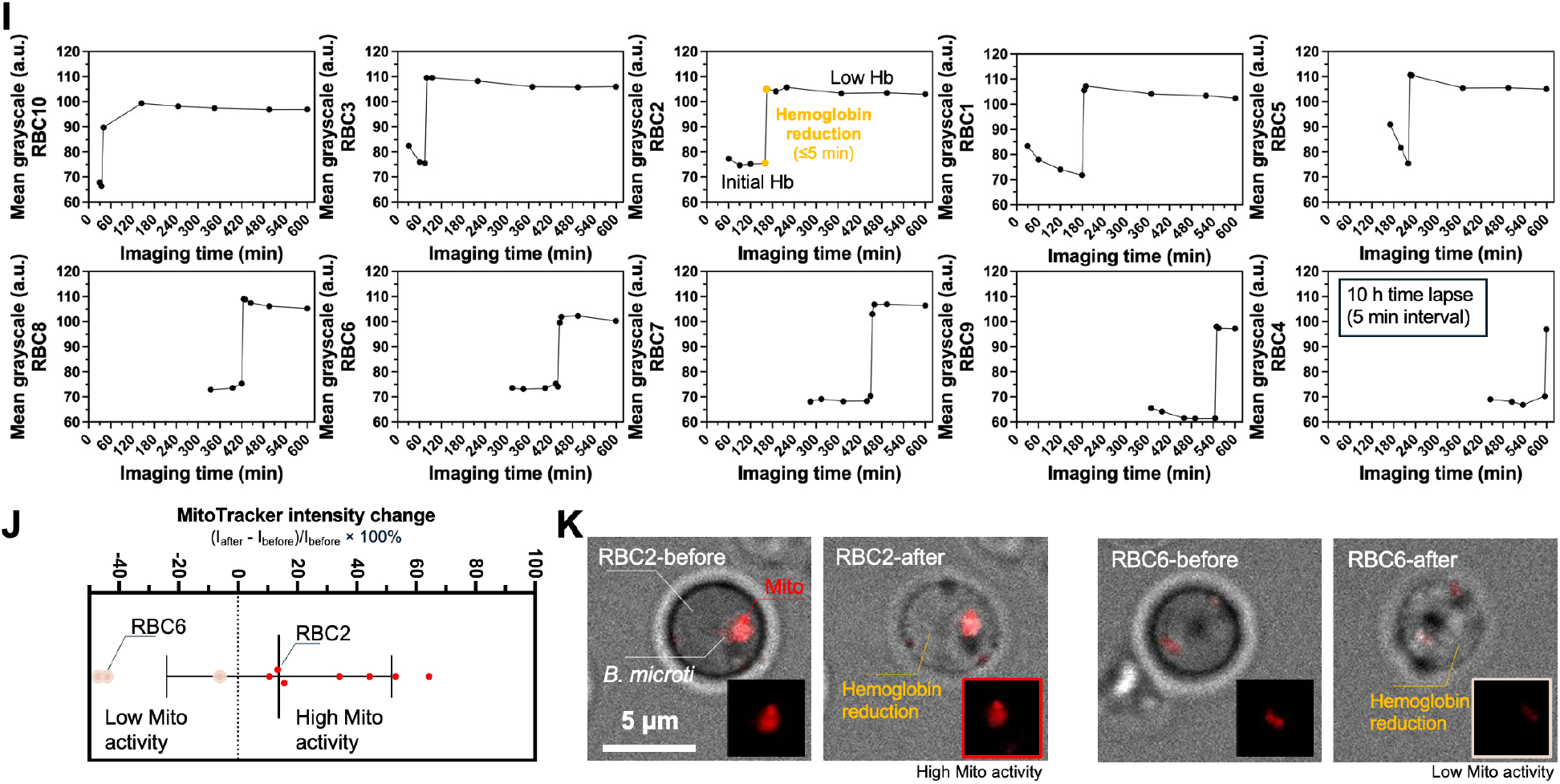
Change of *B. microti*-infected RBCs in whole blood *ex vivo*. (**A**) Composite image [phase + MitoTracker Orange, 8-bit, objective – 63× (oil immersion)] shows the RBCs in the under-oil microchannel (denoted by the white dashed lines) in whole blood at 4 h post blood collection. The white dashed-line boxes highlight the RBCs with initial Hb and low Hb. (**B**) RBCs with initial Hb were identified with a lower grayscale value (i.e., darker cytosol). RBCs with low Hb showed a much lighter cytosol. (**C**) RBC cytosol grayscale normalized to plasma (n = 15) including uninfected (n = 10), infected with initial Hb (n = 20), and infected with low Hb (n = 10). (**D**) Comparison of RBC cytosol grayscale normalized to plasma between partially lysed RBCs in lysis buffer (n = 20) and *B. microti*-infected RBCs with low Hb (n = 10). (**E**) Size (equivalent diameter) and (**F**) Circularity comparison between the RBCs with initial Hb and low Hb. Each data point represents an RBC in its group. RBCs with initial Hb (n = 10); RBCs with low Hb (n = 10). (**G**) Change of the RBC ratio with initial Hb [RBCs with initial Hb/total RBCs) × 100%] through 4 days on μ-Blood (37 °C in air, replicates ×3) (Fig. S6A). The “×” mark on the graph shows the reference point from uninfected blood at 48 h (Fig. 1F). Error bars, mean ± s.d. **P ≤ 0.01, ***P ≤ 0.001, ****P ≤ 0.0001, and ns – not significant. (**H**) Representative image shows the ratio between the RBCs with initial Hb (<10%) and low Hb in whole blood at 48 h post blood collection. (**I**) Hb reduction of *B. microti*-infected RBCs (n = 10) recorded in a time lapse (10 h, 5 min interval, Movie S2). (**J**) Mitochondrial activity (n = 10) probed by MitoTracker Orange (Fig. 3 A and B) before and after Hb reduction shown in (**I**) in a 5-min interval. MitoTracker intensity change = [I_after_-I_before_]/I_before_ × 100%. (**K**) Composite images (phase + MitoTracker Orange) of RBC2 (high Mito activity) and RBC6 (low Mito activity) show the representative mitochondrial activity change before and after Hb reduction.

Compared to the infected RBCs with initial Hb (n = 20), the infected RBCs with low Hb (n = 10) exhibited a significantly higher grayscale value (∼10%, normalized to plasma, P ≤ 0.0001, t-test) therefore a lighter cytosol on phase contrast (Fig. 5C). Compared to uninfected RBCs (n = 10), the infected RBCs with initial Hb (n = 10) also showed a significantly higher grayscale value (∼10%, normalized to plasma, P ≤ 0.0001, t-test) therefore a lighter cytosol on phase contrast. This grayscale difference indicated that *B. microti*-infected RBCs have reduced Hb levels (Fig. 5C). We further compared the RBC cytosol grayscale between partially lysed RBCs in lysis buffer treatment (n = 20) and infected RBCs with low Hb (n = 10) (Fig. 5D). The results showed that the Hb level of infected RBCs with low Hb was significantly higher (∼2% of grayscale, normalized to plasma, P = 0.0008, t-test) therefore slightly darker cytosol than partially lysed RBCs. In addition, the infected RBCs with low Hb (n = 10, ∼4.5 to 6 μm in equivalent diameter) were significantly smaller than the infected RBCs with initial Hb (n = 10, ∼7 to 9 μm in equivalent diameter, P ≤ 0.0001, t-test) (Fig. 5E), both taking largely circular cell shape (average circularity >0.93) (Fig. 5F). The cytoplasmic membrane of infected RBCs with low Hb was clearly identified, indicating a relatively integrated cytoplasmic membrane structure (Fig. 5 B, H, and K). These results together indicate that *B. microti*-infected RBCs can significantly lose Hb content compared to fresh uninfected RBCs but with a cytoplasmic membrane damage comparable to or milder than partially lysed RBCs treated by lysis buffer.

Through a 4-day time-course of imaging on μ-Blood (Fig. 5 G and H), the results showed that most *B. microti*-infected RBCs with initial Hb (>90%, P = 0.0049, One-way ANOVA) had reduced Hb around 48 h post blood collection, cultured at 37 °C in ambient air. The RBC degradation rate with *B. microti* infection from 24 h to 48 h (>90%, Fig. 5G) was significantly higher compared to uninfected blood (∼22%, Fig. 1 E and F). To determine the potential cause of Hb reduction in *B. microti*-infected RBCs over time, we used four infected blood conditions (including whole blood, whole blood diluted with PBS in 1:1, 1:10 and 1:100 vol. ratios) and compared each in parallel on two μ-Blood devices cultured at 37 °C in ambient air (21% O_2_ and 0.04% CO_2_) versus blood physiology with 5% O_2_ and 5% CO_2_ (Fig. S5). On the 1:100 PBS-diluted blood sample (that provided the lowest cell density on surface), we performed a 10 h (5 min interval, 37 °C/air) time lapse to monitor the process of Hb reduction in real time (Fig. 5I). The time lapse showed that Hb reduction occurred rapidly within 5 min (Movie S2), leading to reduced Hb level and still relatively intact RBC cytoplasmic membrane. A recent study reported a type of protein secreted by intraerythrocytic *B. microti* called perforin-like-protein 1 (PLP-1) (*34*), associated with egress of the parasites from an infected RBC. The Hb reduction observed on μ-Blood is likely connected to perforation of the cytoplasmic membrane of infected RBCs as the parasite consumes various components of the RBC membrane for nutrients (*35*). While Hb was rapidly reduced in infected cells an integrated cytoplasmic membrane structure remained intact (Fig. 5K). Additionally, MitoTracker Orange revealed that ∼70% of the parasites retained high mitochondrial activity after Hb reduction (Fig. 5 J and K). This result explains the variation of mitochondrial activity observed at 48 h in infected whole blood (Fig. 5H).

Comparison between the PBS-diluted groups with the raw whole blood samples (Fig. S5) showed that dilution slowed down the process of Hb reduction, reflected by the ratio of RBCs with initial Hb in reference to total RBCs. This indicates a possible connection between Hb reduction and the environmental/signaling factors in plasma. Further, the culture condition of 5% O_2_ and 5% CO_2_ led to an increased Hb reduction ratio (>50%) (Fig. S5F) compared to the ambient air (21% O_2_ and 0.04% CO_2_) culture condition (∼30%) (Fig. S5C). In previous studies we reported that whole blood can retain the physiological level of oxygen (e.g., 5% O_2_ of venous blood) through 2 days in air *ex vivo* (*18*). The dramatic change in oxygen level in whole blood at around 48 h *ex vivo* may act as an environmental or signaling factor from plasma that correlates to the sudden increase of Hb reduction (Fig. 5G, Fig. S6). In addition, the significant difference in the Hb reduction ratio between 5% CO_2_ and air (0.04% CO_2_) (Fig. S5) indicates the pH level of plasma might be another environmental or signaling factor related to *B. microti*-directed damage of infected RBCs.

## DISCUSSION

The development of *in vitro* infection models that preserve the original whole blood microenvironment are critical for advancing our understanding of blood-borne infectious diseases like babesiosis. The μ-Blood platform allows for direct, unprocessed whole blood assays that preserve the physiological conditions of the parasite’s infection microenvironment. Compared to other blood assay platforms established upon closed-system microfluidics (*36*), μ-Blood offers free physical access to samples on the device with minimized system disturbances, preserves a host-specific assay microenvironment for an extended period of time, greatly increasing the potential for studying infection dynamics with difficult-to-culture pathogens, such as *Plasmodium vivax, Enterocytozoon bieneusi*, and *Pneumocystis jirovecii*.

In this study, we demonstrated the capacity of μ-Blood to monitor *B. microti* infection dynamics with whole blood from infected host samples, enabling quantitative, days-long analysis of parasite behavior and parasite-host cell interactions *ex vivo*. Key demonstrations of the assay capacity include label-free identification of *B. microti* in whole blood, improved accuracy for measuring parasitemia levels, and tracking of *B. microti*-directed damage of infected RBCs. μ-Blood revealed critical *B. microti*-RBC interaction highlighted with Hb reduction of infected RBCs and the influence of environmental factors (including blood dilution, blood culture conditions) on Hb reduction dynamics. Future studies on the molecular mechanisms governing *B. microti*-RBC interactions should focus on identifying the factors responsible for RBC cytoplasmic membrane damage, which could reveal new preventive and therapeutic targets.

### Limitations of the method

The current method in this study was demonstrated with standard confocal microscopy configuration, limited to two μ-Blood devices in maximum (16 microchannels ×2) each run (Fig. S1C-III). Sample finding and ROI selection were all performed manually by human operators, limiting data acquisition speed and throughput. To increase the assay efficiency, μ-Blood – enabled by its open microfluidic configuration – can be readily integrated with an automated multiplexed imaging system and robotic device operation platform (e.g., Leica Cell DIVE and BioAssemblyBot 200). Similarly, data processing and analysis in the current method were done manually for proof of principle with limited throughput. Moving forward, we will train AI models (e.g., Leica Aivia AI image analysis software) for automatic cell segmentation, phenotyping, tracking, and statistics report generation. Together, automated device operation, imaging, and AI-powered image analysis will significantly enhance the speed and throughput of the μ-Blood assay.

By preserving the integrity of the host-specific whole blood microenvironment, μ-Blood offers an *ex vivo* solution to the traditional lab animal-centered studies of infection dynamics. Furthermore, the compatibility of μ-Blood with a wide range of optical modalities – including phase contrast, epifluorescence/confocal, and label-free microspectroscopy (e.g., Raman, IR) – enhances its utility for diverse experimental applications. The platform’s impact is likely to be highest in areas where precise extended assays are needed to study complex blood-borne parasite-host cell interactions. Specifically, μ-Blood will facilitate screening of the *in vivo* variables that govern the cultivability of various difficult-to-culture pathogens, which mitigates or removes the heavy reliance on lab animal models in infectious disease studies.

## MATERIALS AND METHODS

### SCID mouse blood collection

Fox Chase SCID mice were purchased from Charles River Laboratories. Female mice (n = 4) were injected with 100 μL of *B. microti* infected blood (LabS1 strain). Blood was collected from each mouse via the submandibular vein 14 days post infection and screened for *B. microti* by viewing Wright-Giemsa-stained blood smears and measuring parasitemia levels using bright field microscopy (Table S1). For confocal imaging, blood samples from uninfected and infected SCID mice were collected and placed in individual tubes containing 15% ACD solution (C3821, Sigma Aldrich) to minimize coagulation. All animal procedures were completed in accordance with the guidelines approved by the University of Pittsburgh Institutional Animal Care and Use Committee (Protocol No. 22051147).

### Parasitemia measurement using traditional blood smear slides

One step Wright-Giemsa stain from Hardy Diagnostics (Santa Maria, CA) was used to stain blood smear slides to visualize *B. microti-*infected RBCs under the microscope (Fig. 2F). Thin blood smears were stained using the Dip procedure following manufacturer’s protocol using Coplin jars. Slides were viewed with oil-immersion 100× magnification using bright field microscopy (S81717, Fisher Science Education Microscope). The average parasitemia level per individual mouse was determined by calculating the number of infected RBCs per 100 RBCs in three different areas of the blood smear slide.

### μ-Blood device fabrication and preparation

#### Polydimethylsiloxane (PDMS) silane-grafted surface

Chambered coverglass [Nunc Lab-Tek-II, 4 well (155382), #1.5 borosilicate glass bottom, 0.13 to 0.17 mm thick, Thermo Fisher Scientific] (Fig. S1C-I) was treated first with O_2_ plasma (Diener Zepto) at 100 W for 3 min and then moved to a vacuum desiccator (Bel-Art F420100000/EMD, S43283, Thermo Fisher Scientific) for chemical vapor deposition (CVD). PDMS-silane (1,3-dichlorotetramethylsiloxane, SID3372.0, Gelest) (20 μL × 2 per treatment) was vaporized under vacuum pumping for 3 min and then condensed onto the substrate under vacuum at RT for 40 min.

The PDMS-grafted surface was thoroughly rinsed with ethanol (anhydrous, 99.5%), deionized (DI) water, and then dried with nitrogen before use.

#### Fabrication of PDMS elastomer stamps

Photomasks were designed in Adobe Illustrator and then sent to a service (Fineline Imaging) for printing. Standard photolithography was used to make a silicon master template that contained all microchannel features (Fig. S1C-II; See (*21*) for further details about the photolithography process). PDMS elastomer stamps were made by pouring a degassed (about 20 min using a vacuum desiccator) silicone elastomer precursor and curing agent mix (SYLGARD 184, Silicone Elastomer Kit, 04019862, Dow Corning) in 10:1 mass ratio onto the master and curing on a hotplate at 80 °C for 4 h. The PDMS elastomer stamps were removed from the master with tweezers and holes were made at the inlet and outlet of a microchannel (Miltex Biopsy Punch with Plunger, 15110, Ted Pella) for the following O_2_ plasma surface patterning.

#### O_2_ plasma surface patterning

The PDMS silane-grafted chamber coverglass was masked by a punched PDMS elastomer stamp and then treated with O_2_ plasma at 100 W for 1 min (Fig. S1C-III). After surface patterning, the PDMS elastomer stamp was removed by tweezers and stored in a clean container (e.g., Petri dish) for reuse.

#### Under-oil microchannels and sample loading

The chemically patterned chambered coverglass was overlaid with 500 μL silicone oil (5 cSt, 317667, Sigma Aldrich) for each well on the 4-well chambered coverglass. Whole blood was distributed onto the microchannels by under-oil sweep distribution (Fig. S1C-III). Briefly, we collected 10 to 20 μL of whole blood in a 1-200 μL large orifice pipette tip (02-707-134, Thermo Fisher Scientific) and then dragged the hanging drop at the end of the tip through the patterned surface. Blood was spontaneously distributed onto the O_2_ plasma-treated areas only. After sweep distribution, we added 500 μL Fluorinert FC-40 (1.85 g/mL at 25 °C, F9755, Sigma Aldrich) to each well on the 4-well chambered coverglass directly into the silicone oil (5 cSt, 0.91 g/mL at 25 °C). Due to the high density of fluorinated oil and immiscibility with silicone oil, FC-40 spontaneously replaces silicone oil and pushes silicone oil to the top layer. Silicone oil was then removed using a pipette placed in the four corners in a well. Fluorinated oil enables minimized media loss via evaporation under oil and extended (days-long) whole blood assay *ex vivo*. In addition, our previous UPLC-MassSpec studies showed no detectable molecule loss (including lipophilic molecules such as short chain fatty acids) from the aqueous phase to the oil phase potentially concerned with liquid-liquid extraction (*24*).

### Confocal microscopy

The imaging of μ-Blood was performed on Leica STELLARIS 5 confocal microscope (Fig. S1A). The objectives used in this study include HC PL APO CS2 20×/0.75 DRY [working distance (WD, 0.62 mm)], HC FLUOTAR L VISIR APO CS2 25×/0.95 WATER (WD, 2.40 mm), and HC PL APO CS2 63×/1.40 OIL (WD, 0.14 mm). The pinhole was set to 1 airy unit (AU) for all imaging. OKOLab onstage incubator was used to control the cell culture environment on the microscope for 37 °C, ambient air or 5% O_2_, 5% CO_2_, and 95% relative humidity (RH). MitoTracker Green (9074, Cell Signaling Technology) is a hydrophobic, membrane-permeable, cationic dye that gets enriched at and covalently binds to the inner membrane of mitochondria (both live and dead). MitoTracker Orange CMTMRos (2252, Lumiprobe) is a hydrophobic, membrane-permeable, cationic dye that stains the inner membrane of live mitochondria. Dead mitochondria lose the fluorescence of MitoTracker Orange over time due to the diminished inner membrane potential.

### Image analysis

Image analyses were completed in Fiji ImageJ as follows i) Cell count was performed using “Plugins – Analyze – Cell Counter”. ii) Singal (e.g., grayscale, fluorescence) intensity and morphology (including size and shape) analyses were performed using “Analyze – Measure”. “Set Measurements” for intensity analysis included “Mean gray value, Standard deviation, and Min & max gray value”. Set Measurements for morphology analysis included “Shape descriptors and Feret’s diameter”. The ROI was defined using the shape selection tools on the objects of interest. Supporting movies were edited in DaVinci Resolve 19.

### Data visualization and statistical analysis

Raw data was directly used in statistical analysis with no data excluded. Data was averaged from at least 3 replicates and presented as mean ± standard deviation (s.d.) if applicable. Data plotting and visualization were performed using GraphPad Prism (Ver. 10.4.1), Microsoft Excel, and Microsoft PowerPoint. All statistical analyses were performed using t-test (for paired data) or One-way ANOVA (for comparison of multiple data).

## Author contributions

T.H.R.N., D.M.T., and C.L. conceived this project with equal contribution. C.L. developed the μ-Blood platform, designed and performed the experiments, and performed data analysis/visualization (at Carnegie Mellon University). E.G.B. and D.M.T. collected blood samples from *B. microti* infected mouse colony maintained at the mouse facility (at the University of Pittsburgh) and delivered the blood samples between labs. E.G.B. and A.L.A. performed blood smear characterization on parasitemia level of the blood samples (at the University of Pittsburgh). T.H.R.N. and D.M.T. supervised this project. C.L. wrote the manuscript and all authors revised it.

## Data and Materials Availability

All data needed to evaluate the conclusions in the paper are present in the paper and/or the Supplementary Materials.

