## Supporting Information for "*In vitro* monitoring of *Babesia microti* infection dynamics in whole blood microenvironments"

**This SI file includes:**

Figures S1 to S6

Table S1

Movies S1 and S2

A

B

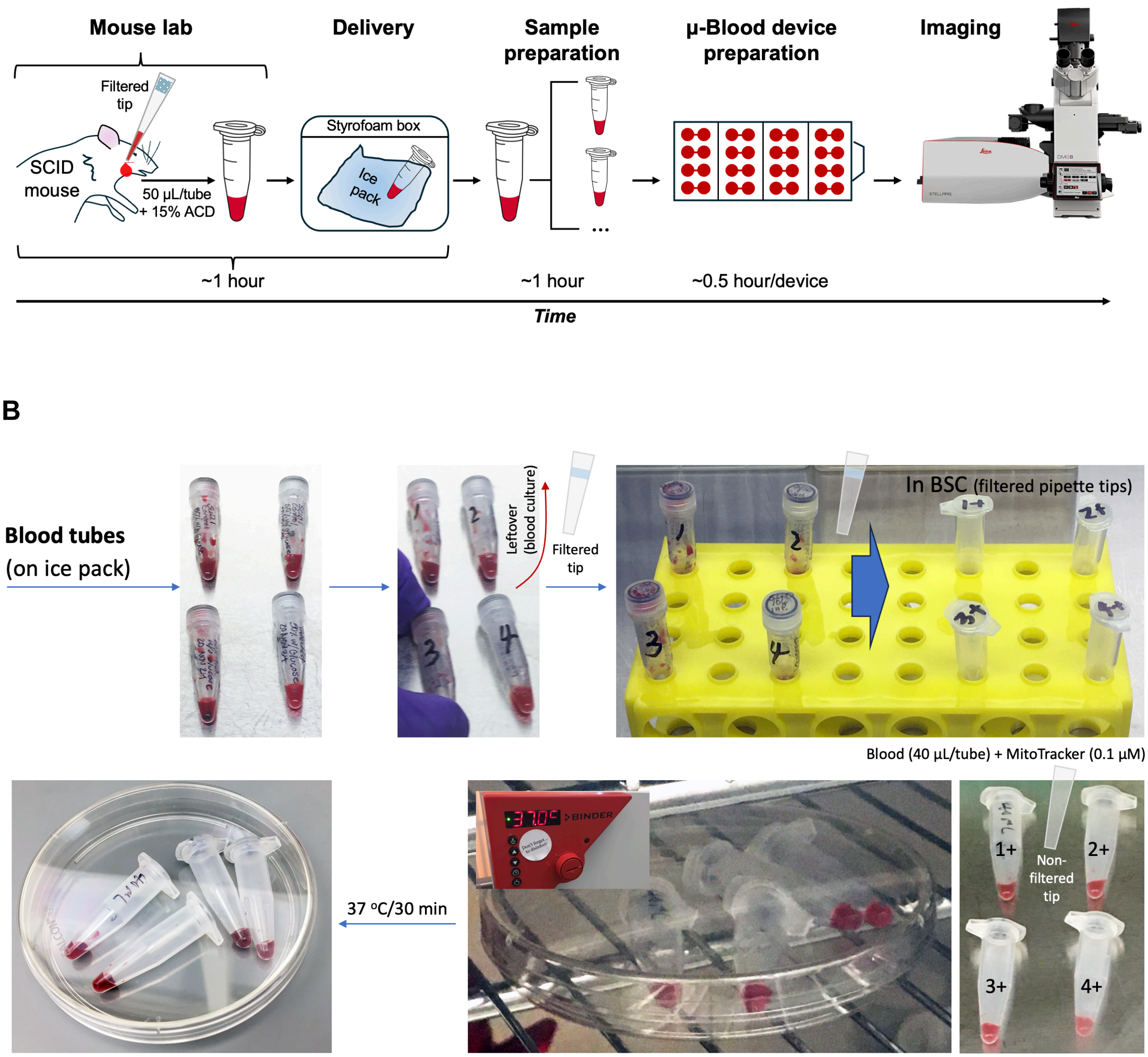
Mouse lab
Delivery
Sample preparation
μ-Blood device preparation
Imaging
Filtered tip
SCID mouse
50 μL/tube + 15% ACD
Styrofoam box
Ice pack
~1 hour
~1 hour
~0.5 hour/device
Time
Blood tubes
(on ice pack)
Leftover (blood culture)
Filtered tip
In BSC (filtered pipette tips)
Blood (40 μL/tube) + MitoTracker (0.1 μM)
37 °C/30 min
1+
2+
3+
4+
Non-filtered tip

**C**

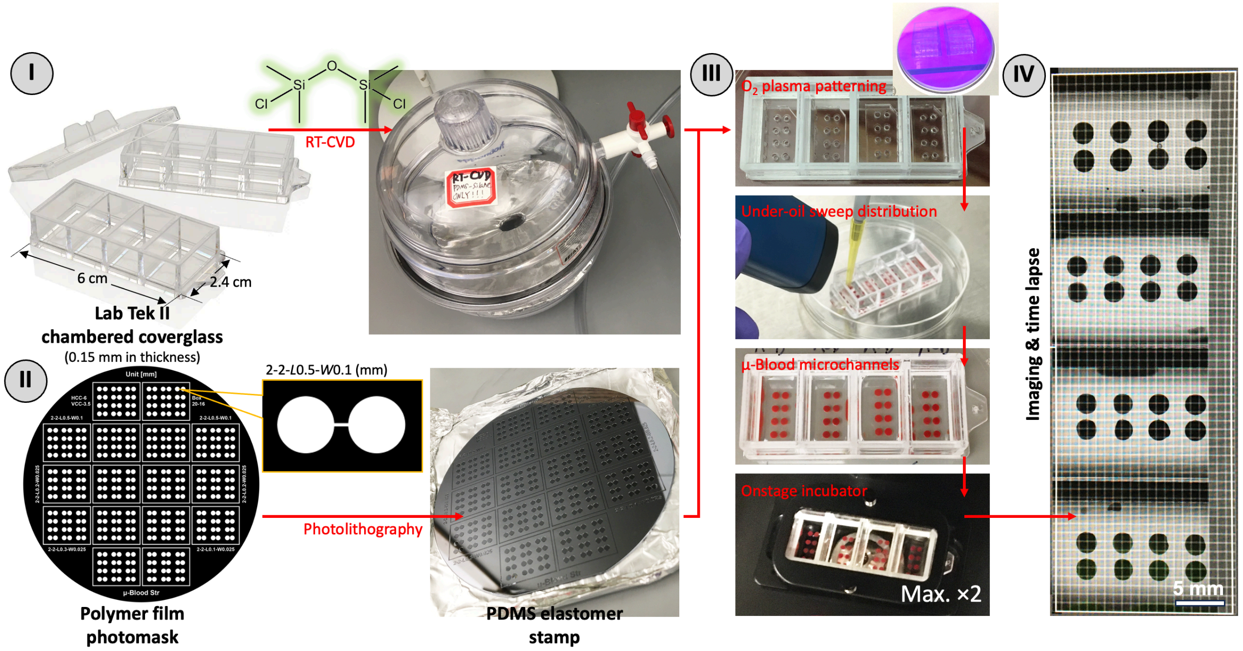


**Fig. S1 Workflow of μ-Blood *Babesia* assay.** (**A**) Average time use in each step from blood collection to data collection. (**B**) Representative blood sample preparation before sample loading on the device. μ-Blood can handle small blood sample volume for 10+ μL. In each experiment, the leftover blood was put in culture with LB media as bacterial contamination control. (**C**) μ-Blood device fabrication and operation including I – PDMS silane CVD (done independently before the assay), II – PDMS stamp preparation (done independently before the assay, stamps reusable), III – surface patterning and sample loading (maximum of two μ-Blood devices at a time on microscope), and IV – imaging and time lapse (a representative preview of the entire device, rotated 90° counterclockwise). μ-Blood device preparation in (**A**) corresponds to III in (**C**). Imaging in (**A**) corresponds to IV in (**C**).

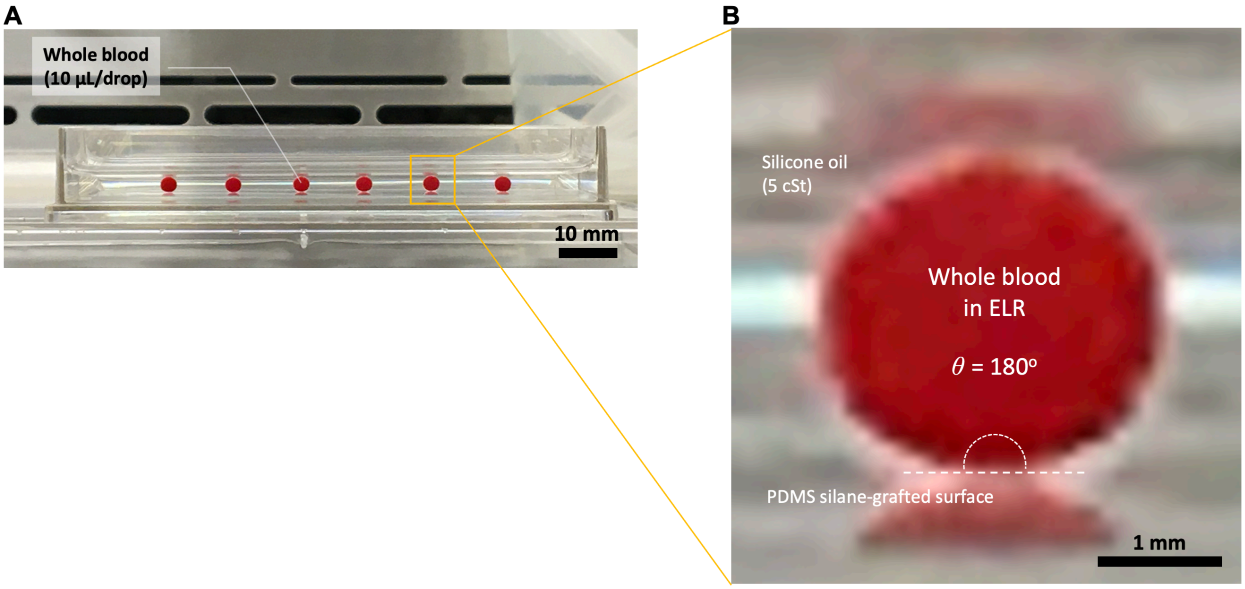


**Fig. S2 ELR of whole blood on PDMS silane-grafted glass surface under oil (silicone oil, 5 cSt).** (**A**) Six drops of whole blood (sheep, defibrinated, 10 µL per drop) in a Nunc OmniTray. (**B**) Zoomed-in picture of ELR blood drop shows the droplet profile with a Young's contact angle $\theta$ = 180º.

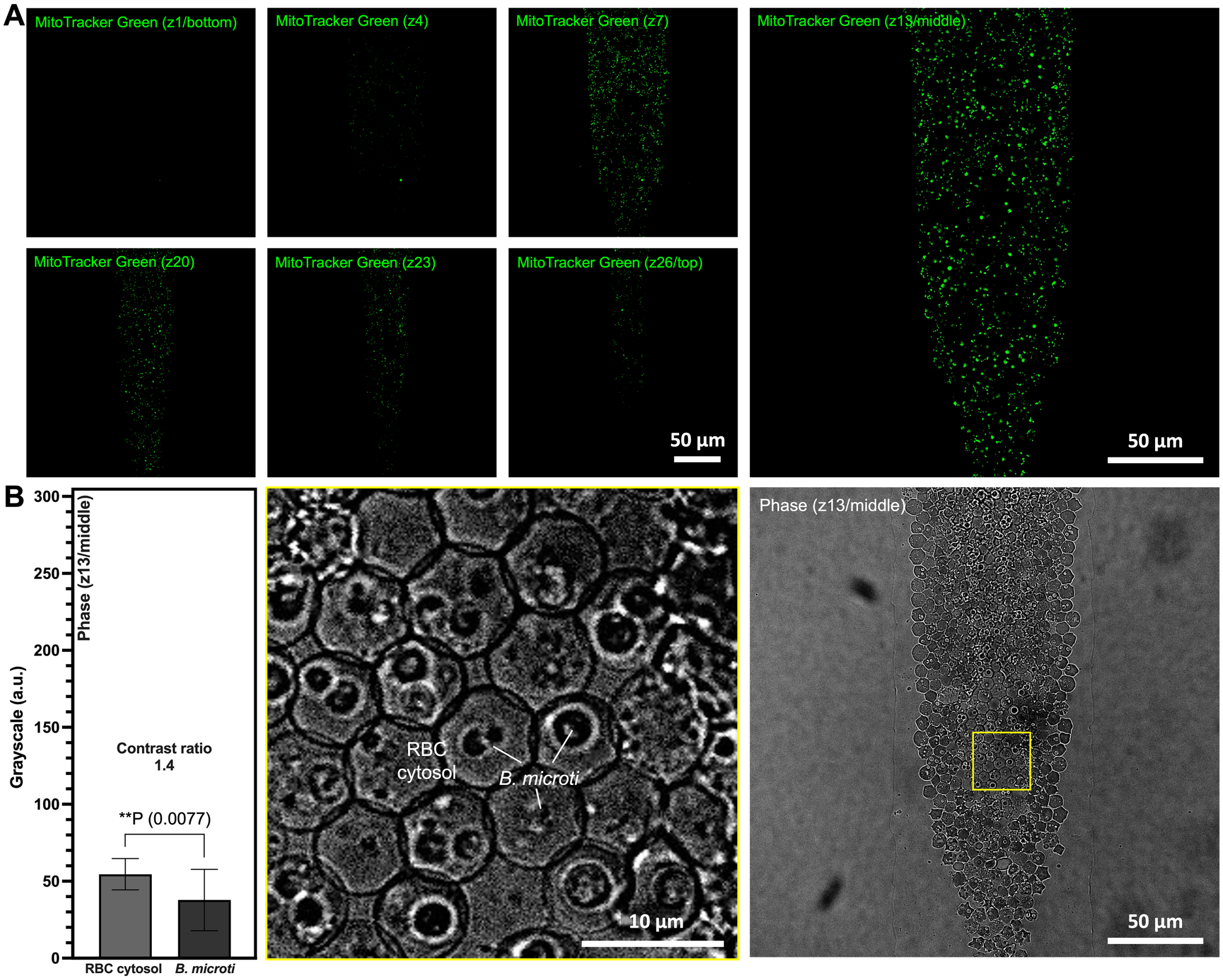
A
MitoTracker Green (z1/bottom)
MitoTracker Green (z4)
MitoTracker Green (z7)
MitoTracker Green (z13/middle)
MitoTracker Green (z20)
MitoTracker Green (z23)
MitoTracker Green (z26/top)
50 µm
50 µm
B
Phase (z13/middle)
Grayscale (a.u.)
300
250
200
150
100
50
0
Contrast ratio
1.4
**P (0.0077)
RBC cytosol
B. microti
RBC
cytosol
B. microti
10 µm
Phase (z13/middle)
50 µm

**C**

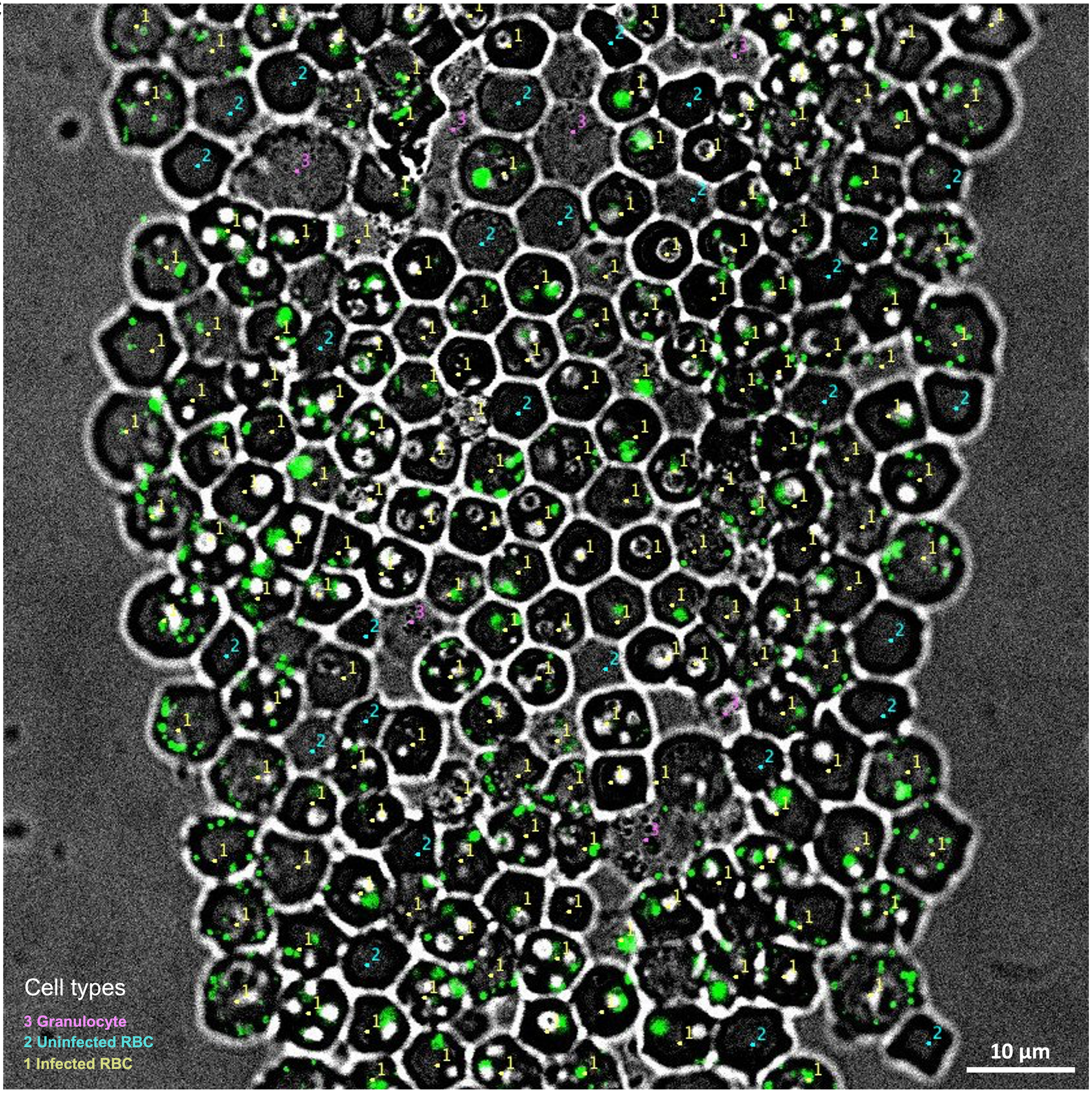


**Fig. S3 Confocal images and z-stack slices of *B. microti*-infected SCID mouse blood on µ-Blood.** (**A**) Selected MitoTracker Green images from z-stack in Fig. 2. The middle slice of MitoTracker Green shows the maximum fluorescence intensity and coverage of mitochondria. (**B**) The middle slice of phase from z-stack shows the minimum contrast ratio (1.4) between RBC cytosol and *B. microti* compared to the bottom (44.6) and top (37.3) slice (Fig. 2E). Error bars, mean ± s.d. $^{**}P \leq 0.01$. (**C**) A representative image of *B. microti*-infected blood with the cell-count markers. Cell types include 1. Infected RBC, 2. Uninfected RBC, and 3. Granulocyte.

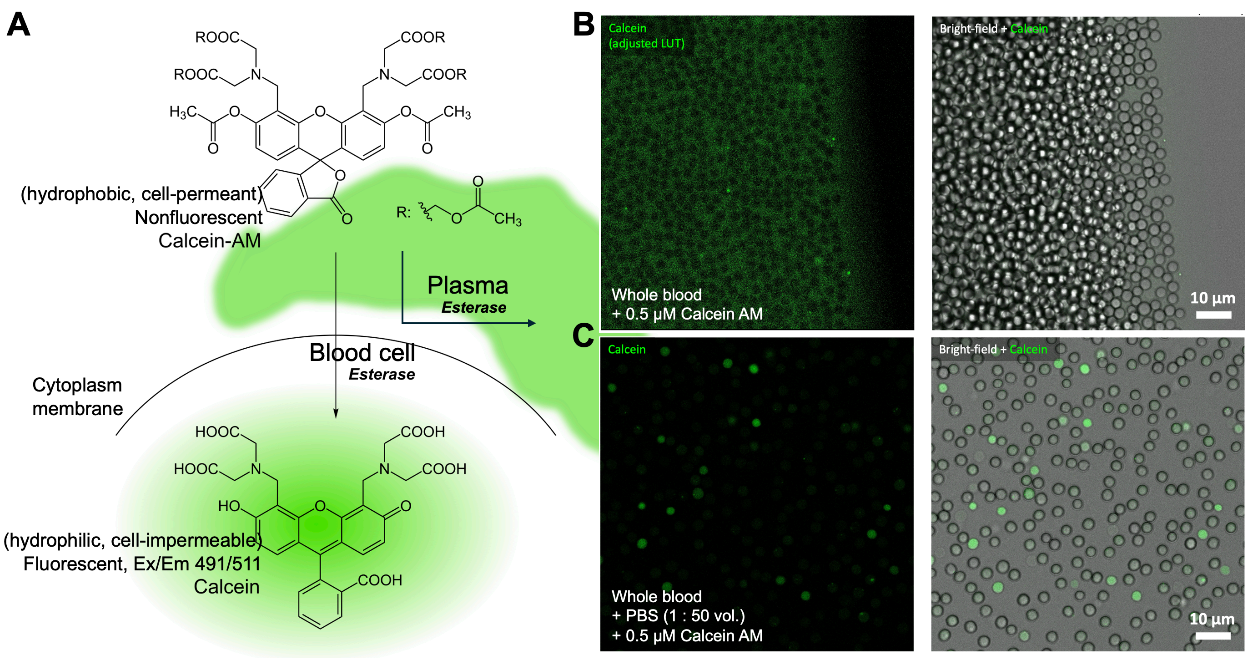


**Fig. S4 Calcein AM cell viability stain in *B.* microti-infected whole blood.** (**A**) Schematic shows the molecular mechanism of Calcein staining. (**B**) High fluorescence background and the flipped contrast between plasma (green/bright) and blood cells (dark) due to the hydrolysis of Calcein AM in plasma. (**C**) Low fluorescence background and the normal contrast between plasma (dark) and blood cells (green/bright) after PBS dilution of whole blood (1:50 vol.).

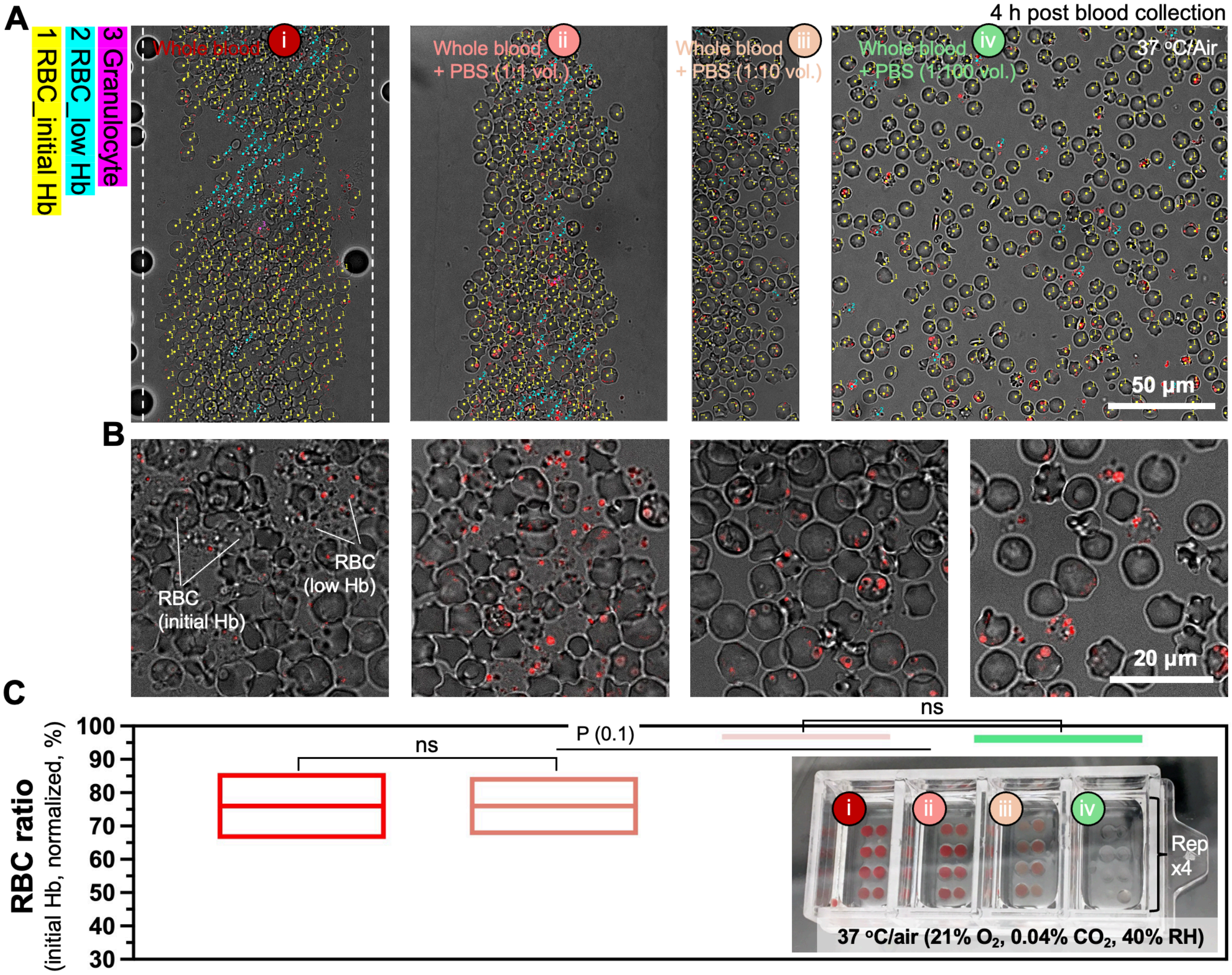
A
4 h post blood collection
1 RBC_initial Hb
2 RBC_low Hb
3 Granulocyte
Whole blood
Whole blood + PBS (1:1 vol.)
Whole blood + PBS (1:10 vol.)
Whole blood + PBS (1:100 vol.)
37 °C/Air
50 μm
B
RBC (initial Hb)
RBC (low Hb)
20 μm
C
RBC ratio
(initial Hb, normalized, %)
100
90
80
70
60
50
40
30
ns
P (0.1)
ns
i
ii
iii
iv
Rep x4
37 °C/air (21% O2, 0.04% CO2, 40% RH)

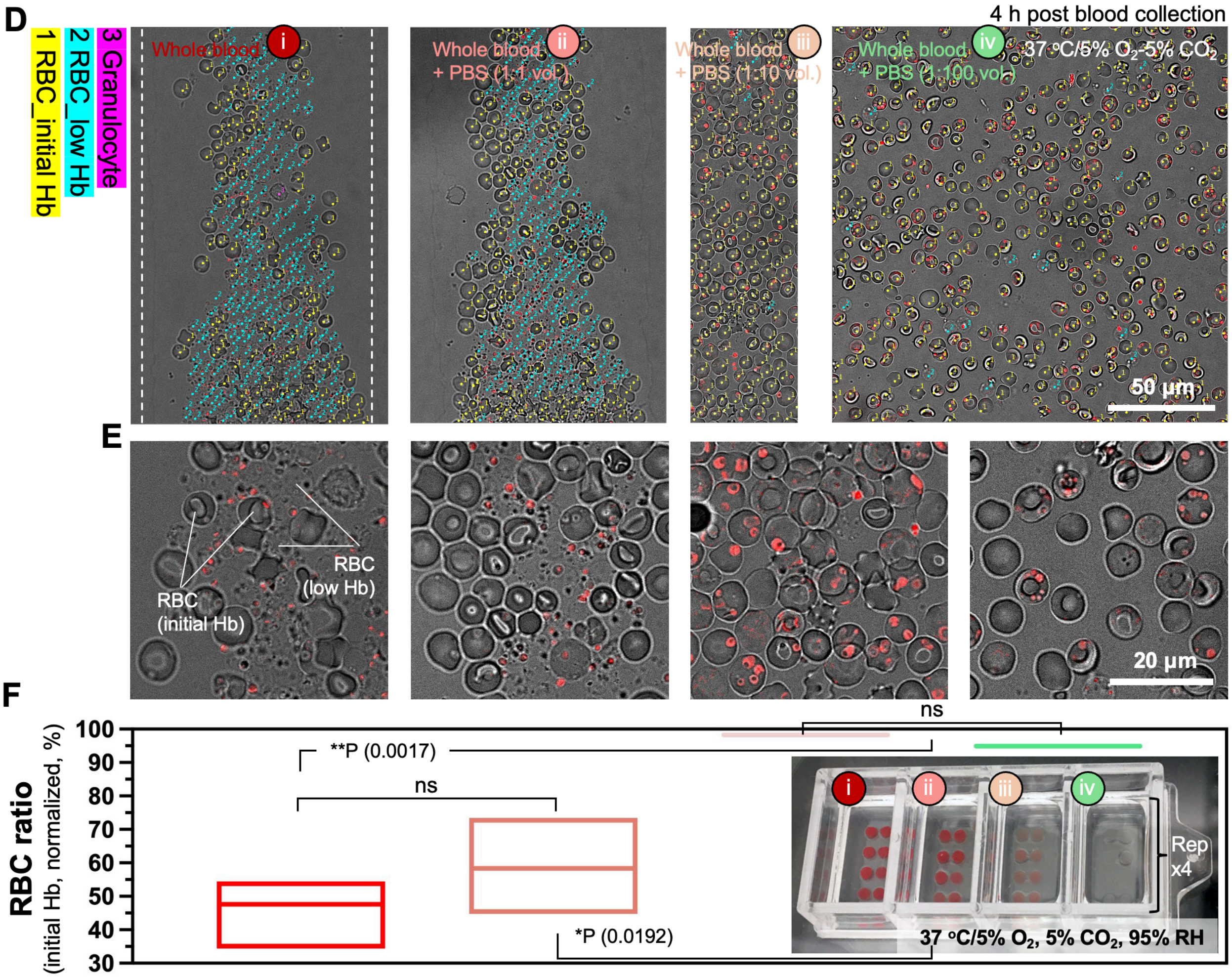


**Fig. S5 Hb reduction of *B. microti*-infected RBCs in different culture environments.** (**A**) Cell counts from four culture conditions including i – whole blood (without PBS dilution), ii – whole blood + PBS (1:1 vol. dilution), iii – whole blood + PBS (1:10 vol. dilution), and iv – whole blood + PBS (1:100 vol. dilution) in ambient air (37 °C, air – 21% $O_2$, 0.04% $CO_2$, ambient <40% RH). The cell count was performed with three cell types including 1. RBC with initial Hb, 2. RBC with low Hb, and 3. Granulocyte. (**B**) Zoomed-in images show the RBC Hb level (grayscale) and *B. microti* (MitoTracker Orange) corresponding to each condition in (**A**). (**C**) Comparison of the ratio of RBCs with initial Hb through the four culture conditions at 4 h post blood collection. (**D**), (**E**), and (**F**) The corresponding results of the four culture conditions with the samples cultured in a $CO_2$ incubator (37 °C, 5% $O_2$, 5% $CO_2$, 95% RH). The whole blood in this test was from the same blood sample collected from a *B. microti*-infected SCID mouse. Insets in (**C**) and (**F**) showed the µ-Blood devices in this test. Error bars, mean ± s.d. *P ≤ 0.05, **P ≤ 0.01, and ns – not significant.

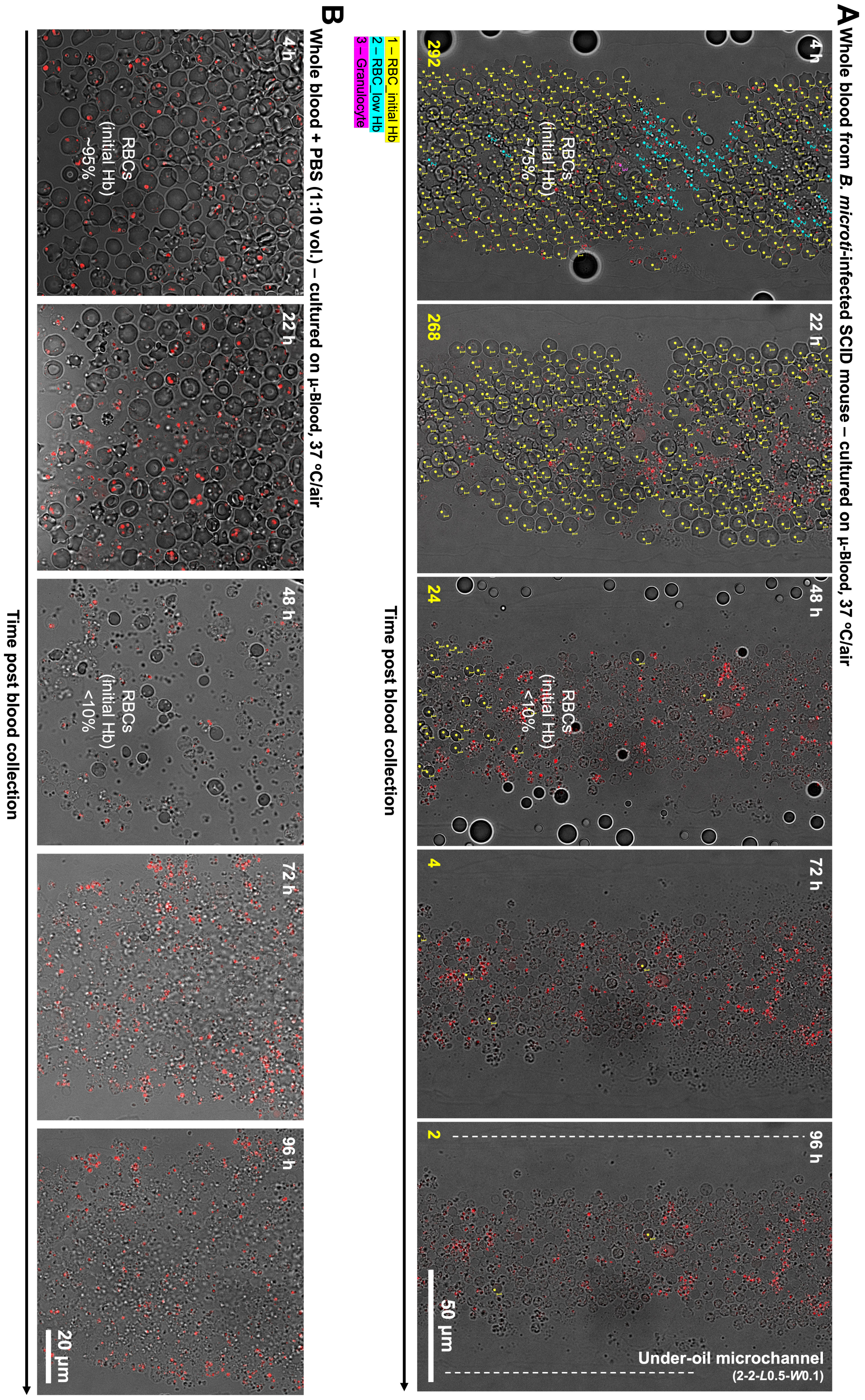


**Fig. S6 Effect of blood dilution on Hb reduction of *B. microti*-infected RBCs.** (**A**) Composite images (phase + MitoTracker Orange) show the change of Hb level in *B. microti*-infected RBCs (yellow numbers) in whole blood (without dilution) through 4 days. (**B**) Composite images (phase + MitoTracker Orange) show the change of Hb level in *B. microti*-infected RBCs in whole blood + PBS (1:10 vol. dilution) through 4 days.

**Table S1 Parasitemia level measured with blood smear and μ-Blood.**

| Mouse ID | Date of blood collection | Parasitemia level (blood smear w Wright-Giemsa stain)* | Parasitemia level (μ-Blood) | Corresponding figure |
|---|---|---|---|---|
| SCID106 | 2024-10-15 | 59.8 ± 3.8% | 82.2 ± 4.6% | Fig. 2 |
| SCID106 | 2024-11-20 | 79.7 ± 5.7% | | Fig. 3 |
| SCID106 | 2024-12-18 | 83.0 ± 2.6% | | Fig. 5 |

*Blood smear comes with limited accuracy and consistency, especially on distinguishing infected RBCs with a single parasite cell from uninfected RBCs (Fig. 2F).

**Movie S1 Phase contrast flip from confocal z-stack imaging.** 5.2 s per frame_ FPS6_31.2× speed_2.25 min z-stack from bottom (z1) to top (z26).

**Movie S2 Hb reduction of a *B. microti*-infected RBC.** 5 min per frame_FPS12_3600× speed_10 h time lapse.
